# A novel feeding mechanism: Sunbirds drink nectar via intralingual suction

**DOI:** 10.1101/2024.05.14.594085

**Authors:** David Cuban, Cynthia Wang-Claypool, Yohanna Yohanna, Colleen T. Downs, Rauri C. K. Bowie, Fabian Brau, Steven D. Johnson, Alejandro Rico-Guevara

## Abstract

Nectarivory has independently evolved many times among birds, yet little is known about the diversity of feeding mechanisms that enable specialized taxa to efficiently collect this energy-rich resource. Multiple avian groups have converged on evolving elongated bills and tube-like tongues adapted for nectar extraction. Old World sunbirds (family Nectariniidae) stand out as having the greatest degree of convergence in bill and tongue morphology with the well-studied and highly-specialized New World hummingbirds (family Trochilidae) which fill their tongues via elastic filling. However, using museum specimens, high-speed video, and fluid modeling, we show that sunbirds use a previously undescribed and unique drinking mechanism not found in any other animal: intralingual suction through the inside of hollow tubular tongues, a remarkable feat for animals without lips or cheeks.

## Introduction

Feeding is a fundamental need of all organisms, and a variety of food acquisition mechanisms have been studied for centuries (*1*–*5*). Although most vertebrates use lapping or licking to drink, a few groups can suck from large pools of liquid by creating a small oral opening connected to an expandable oral cavity (*e*.*g*., some mammals; *6*), which requires their lips to be in contact with the liquid. Only one vertebrate, a species of bat, has been reported to transport liquid along the length of its tongue, using lateral grooves on the surface of their tongues (*7*).

For these bats, however, suction is not possible because the edges of the tongue grooves do not overlap to form a seal, and their tongues are not hollow but rather muscular. For vertebrates without lips or cheeks, suction is an even greater challenge. In birds, suction feeding is restricted to a few taxa (*e*.*g*., filter feeders, pigeons) and it is accomplished with the bill being submerged in water and the tongue acting as a piston (review in *8*). There are no known examples of vertebrates able to move liquids through their tongues using active suction, which involves generation of an internal pressure differential. Nectar-feeding insects (e.g., lepidopterans), and other invertebrates that feed on liquids via suction, have evolved proboscides and other mouthparts analogous but developmentally distinct from tongues and require muscular pumps inside their heads, which are also non-homologous to any vertebrate structure (*9*–*11*). The dietary niche of nectar feeding is a prime candidate for novel feeding mechanisms to be discovered as the animals rely upon efficient consumption of small and hard-to-reach amounts of liquid food to offset the elevated caloric needs of this demanding lifestyle (constant commute among flowers and/or intense resource defense; (constant commute among flowers and/or intense resource defense; *12, 13*).

Vertebrates are responsible for 3-25% of plant pollination across habitats, and amongst them, birds are the most diverse pollinators, with over 920 species pollinating a plethora of plant groups (*14*–*18*). Within birds, specialized nectar feeding has evolved independently in approximately 30 clades (*19, 20*), which exhibit varying degrees of morphological, functional, and behavioral convergence (*20*–*24*). Some of the most identifiable convergent traits to this specialized dietary niche occur in the feeding apparatus. Bill and tongue similarities are evident among taxa in distant geographic regions; for example, hummingbirds (Trochilidae) in the Americas, honeycreepers (Thraupidae) in Hawaii, sunbirds (Nectariniidae) in Africa and Asia, asities (Philepittidae) in Madagascar, and honeyeaters (Meliphagidae) in Australia (*20, 25*–*27*). Of the many nectar-feeding birds across the world, the nectar drinking mechanisms are best known for hummingbirds (*28*–*30*). A taxon morphologically similar to hummingbirds, yet phylogenetically distinct, are the sunbirds whose feeding mechanism has not been previously experimentally investigated (*26*).

There are 142 species of nectarivorous sunbirds, and their ranges span Africa, southern Asia, and the Malay Archipelago (*31, 32*). For over a century, hypotheses about their feeding mechanics have been proposed (*27, 31, 33, 34*), with capillarity as the most recently noted (*35, 36*), yet these ideas have not been tested (*24*). The mechanics of drinking nectar have been explored across a diversity of hummingbird species, and given that both sunbirds and hummingbirds have elongated and narrow bills, housing equally long and distally hollow, bicylindrical tongues (*27, 37*), we expected that these groups would employ similar feeding mechanisms. We studied sunbirds from both Africa and Asia, sampling seven species from multiple clades across the phylogeny, as well as species across a wide range of body and bill sizes. To comprehensively study sunbird feeding mechanics and the structures involved in the process, we used micro-computed tomography (µCT) scans and microscope image analysis, high-speed videos of sunbirds drinking from transparent artificial flowers, kinematic analyses, and fluid dynamics models.

## Results

### Tongue morphology, kinematics, and nectar flow

We used microscope photography and µCT scans of dissected sunbird tongues, including some of the same specimens we filmed, to detail their morphology (Fig. 1, A and B). The sunbird tongue can be described along three distinct sections: (1) distally, the tongue is bifurcated, and made of medially curling, keratinous, and very thin walls that form two hollow cylinders (Fig. 1C, cross section [XS] 6); (2) in the middle section, the tongue is no longer bifurcated, the walls are less curled inward with their edges open toward each other, forming a single elliptical cylinder, hollow across its entire length (Fig. 1C, XS 4-5); (3) in the proximal section, the walls of the lingual cylindrical groove are much thicker, and the tongue opens to form a “U”-shaped channel with its edges flared out laterally (Fig. 1C XS 1-3).

**Fig. 1.**
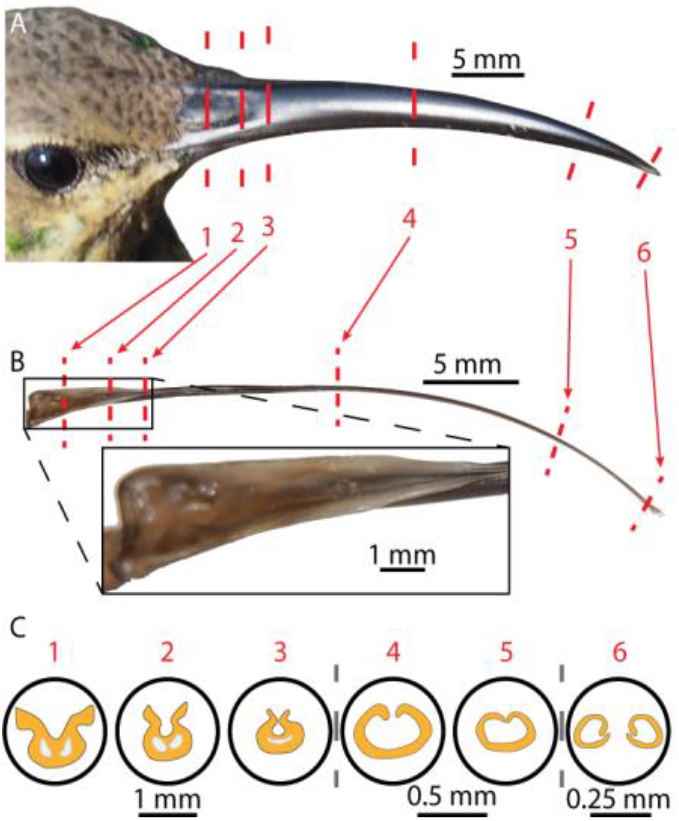
Sunbird bill, tongue, and cross-sections from microCT images. The sunbird tongue can convey fluid along its entire length, with the lateral edges of the tongue base capable of sealing against the inside of the upper bill. **(A)** Dorsolateral view of a *Nectarinia famosa* from the study. Red dashed lines correspond with the locations of the µCT cross-sections. **(B)** Dorsolateral view of the dissected tongue from the above *N. famosa* with cross-section lines marked. **(Inset)** magnified proximal portion of the tongue showing dorsal and lateral edges. **(C)** Enlarged cross-sections from different locations along the length of the sunbird bill and tongue. Cross sections (XS) 1-3 show the proximal portion of the tongue, the “U”-shaped section at the back that becomes narrower further down the tongue.Orange parts are keratin and light gray parts are the internal paraglossal bones located inside the thicker keratin which tapers down to very thin walls towards the distal end. XS 4 and XS 5 show the hollow middle section formed by the inward curling edges of the tongue walls, forming an elliptical space. XS 6 shows the bifurcated tip of the tongue, with the walls rolling inward to form tube-like structures.

To visualize the nectar flow and tongue movement of sunbirds feeding on artificial nectar, we fed birds *ad libitum* with red-dyed (to enhance flow visualization) sucrose solutions of 20% w/w. Dorsal and lateral videos, captured using synchronized high-speed cameras, showed that all sampled sunbirds (nine individuals across seven species, Table S1) fed in similar ways. 1) They protruded their tongue past the open bill tips until the distal portion of the lingual cylinders entered the nectar. 2) They kept their tongue inserted while nectar flowed towards the mouth, filling the internal tongue capacity. 3) They retracted the tongue only partially to swallow, leaving the distal cylinders partly protruded from the bill and filled with nectar. And 4) without squeezing the tongue with the bill tips, they re-extended it for the next lick cycle (*e*.*g*., Video S1). Using the dorsal videos of the transparent nectar chamber, we tracked the tongue motion and fluid flow during the lick cycle for all individuals (e.g., Fig. 2A). Figure 2 shows the data for a malachite sunbird (*Nectarinia famosa*). The position of the tongue tip, nectar meniscus in the reservoir, and nectar meniscus in the tongue are measured relative to a fixed point at the base of the artificial flower (Fig. 2A) over multiple tongue extensions and retractions per feeding bout (*e*.*g*. Fig. 2B). After initial videography experiments, we did not focus on recording or tracking the bill tips as they remained open and relatively stationary during feeding so we determined that, surprisingly, they were not actively involved in the sunbird nectar collection biomechanics (*e*.*g*., Video S1). By subtracting the position of the reservoir meniscus from the tongue tip position, we found the immersion depth to be limited to a narrow band (between 1.3 and 2.1 mm), despite the receding reservoir meniscus and the high tongue reciprocation rate, with an average lick period of ∼0.11 s (Fig. 2C). Notably, sunbirds did not immerse their tongues in the expected way relative to other nectar-feeding birds; e.g., hummingbirds and honeyeaters dip their tongues deep inside the nectar and continuously reciprocate them faster in and out the nectar except for some honeyeaters (*38, 39*). Sunbirds immersed only the very distal portion of their tongue tips in the nectar and kept them stationary at the point of maximum protrusion, performing long “pauses”, which comprised roughly half of the entire lick cycle. We subtracted the reservoir meniscus position from the tongue meniscus position to calculate the nectar column position in the tongue relative to the nectar reservoir, *h* (Fig. 2D). Each of the licks were overlaid with each other relative to the tongue insertion time comparing the position, *h*, of the nectar meniscus in the tongue (Fig. 2E). We averaged all the tongue meniscus positions for each lick, at the same relative time in the lick cycle, and found a linear relationship between meniscus position and time (Fig. 2F).

**Fig. 2.**
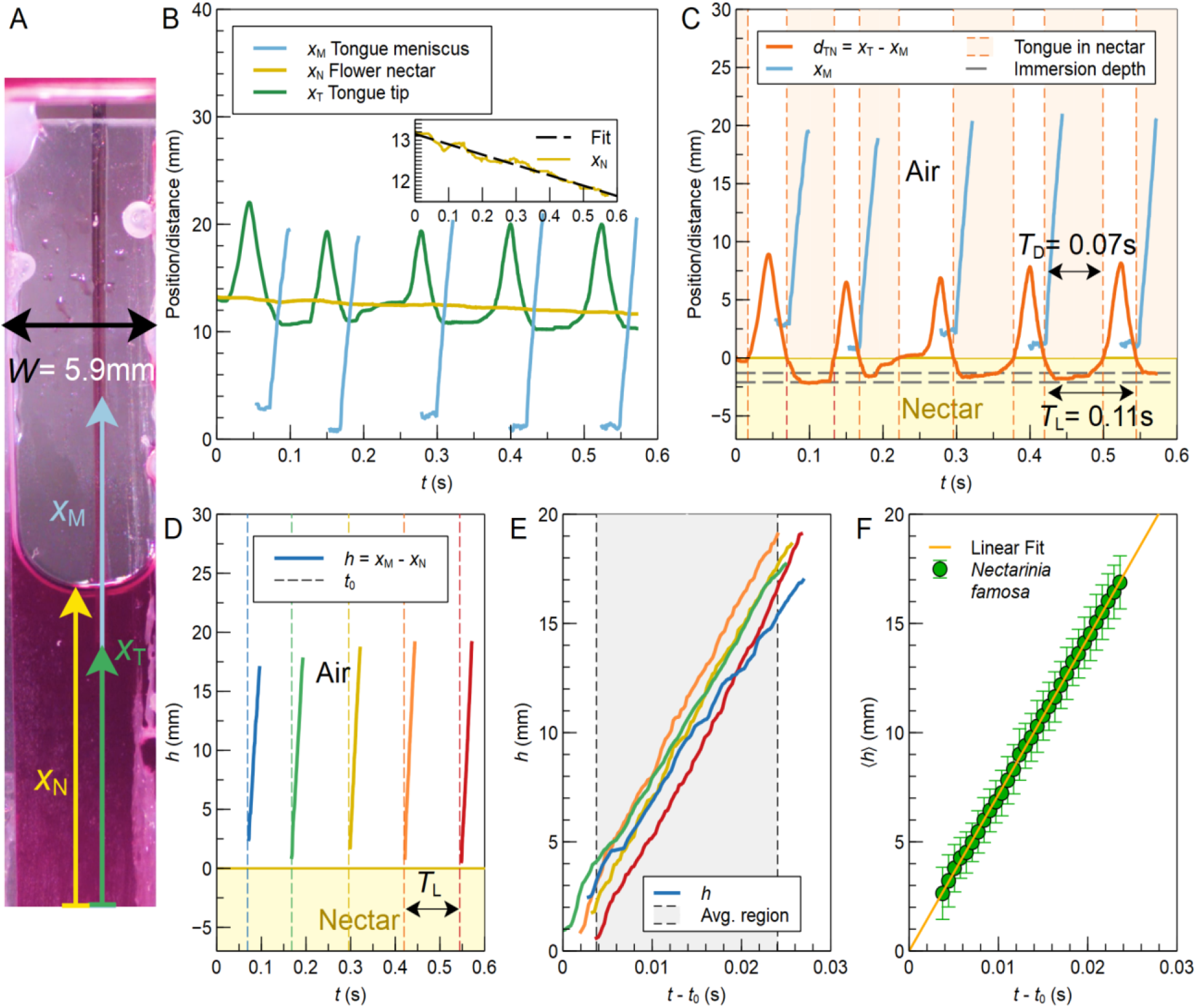
Kinematic measurements of a *Nectarinia famosa* tongue and red colored nectar during multiple lick cycles. **(A)** Video frame showing the nectar reservoir and the tongue tips immersed in the nectar (dyed magenta to improve visualization). Arrows indicate the position of the nectar meniscus of the reservoir, x_N_, the meniscus inside the tongue, x_M_, and the tongue tip, x_T_, with respect to an arbitrary fixed point. **(B)** Position of x_N_, x_T_ and x_M_ as a function of time. **Inset**: Position of x_N_ as a function of time together with a linear fit. **(C)** Position of the tongue tip relative to the nectar reservoir, dTN, as a function of time. Tongue insertion depth varies between 1.3 and 2.1 mm. **(D)** Nectar position, h, in the tongue relative to the nectar reservoir as a function of time. t_0_ corresponds to the time at which the tip of the tongue touches the nectar in the extension phase for each capture process. **(E)** Position of h as a function of t − t_0_ for 5 licks and the time interval used to average h values. **(F)** Average and linear fit of h data in panel E.

### Fluid dynamic modeling

We compare the nectar meniscus rise in the tongue averaged across multiple licks (e.g., Fig 2) among individuals from each species (Table S1) with three feeding mechanics fluid models of increasing complexity. To inform the models (Table S2), we collected all of the specimens for which we obtained feeding performance data, and measured their tongues via dissections and µCT scans (e.g., Figs. 1, S2). First, as the one that has been favored recently (*35, 36*), we present the expectations from the capillarity model, accounting for the surface tension of the sucrose solution at the air-nectar interface inside the tongue and the viscosity of the nectar. As the model shows significant deviation relative to the measured meniscus position, especially after a few initial milliseconds, capillarity on its own does not explain our observations (Fig. 3). Second, also modeling capillarity, but including the inertial effects of accelerating the fluid from a static position, brings the initial meniscus position closer to that of the observed values, especially during the initial fluid climb (first few milliseconds), after which the empirical data still mismatch (Fig. 3). Third, we included a time-varying applied difference of pressure in the model, which resulted in a satisfactory fit of the output with our empirical measurements, and provided support for active suction being the primary mechanism at play (Fig. 3). See “Derivation of fluid models” in the Supplementary Materials for detailed derivations of the fluid models.

**Fig. 3.**
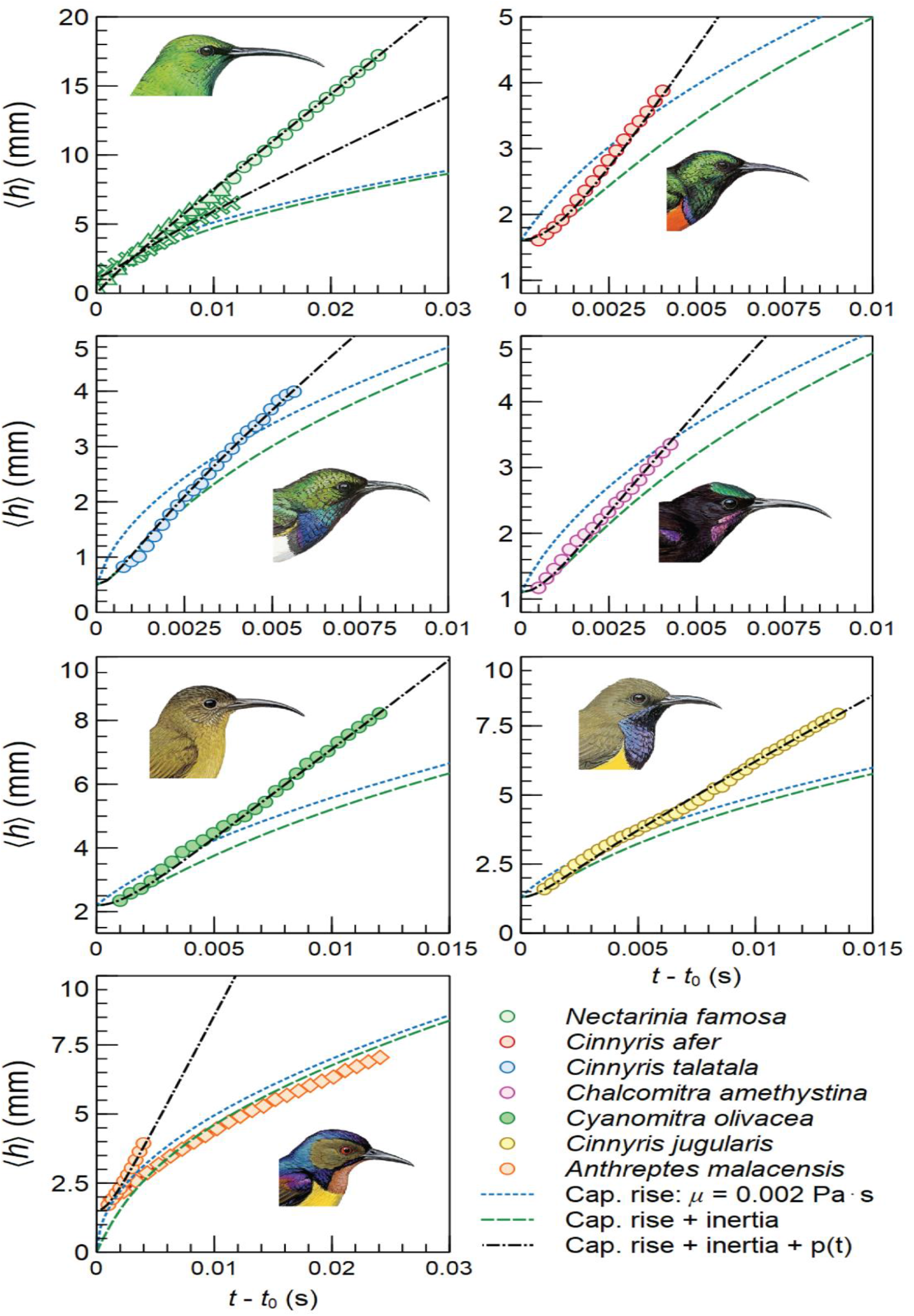
Intralingual suction across sunbird species. Change of ⟨h⟩, nectar position in tongue, as a function of time together with the change predicted by capillary rise only (blue dotted curve), by capillary rise and inertial effects (green dashed curve) and by capillary rise, inertial, effects, and a difference of pressure across the tongue’s length varying linearly in time (black solid curve). These predictive models are adjusted based on the tongue morphology for each species. Data averaged across multiple licks from individuals from each species in the study are plotted (see Table S1). Differences in lick sequence duration (video frames) are due to different tongue lengths and bill tip to nectar reservoir distances. We only observed nectar distance motion typical of capillarity alone in one foraging bout of *Anthreptes malacensis*, and the same individual fed in the more common mode in another video where the tongue was much less extended (both plotted).

The slope, or velocity, of the nectar meniscus in the birds’ tongues matched the predicted slope of a pressure-dominated flow, especially during the portions of the tongue filling process where capillarity is too slow to account for the observed flow. The applied difference of pressure itself is found to increase linearly with time, implying that whatever internal motion the bird is doing to generate suction must be happening continuously during the period of linear advance of the intralingual nectar meniscus. One individual, the brown-throated sunbird *Anthreptes malacensis* (the species with the shortest bill and tongue in the study, Table S1), showed evidence of using suction during one of its feeding bouts, and of a second mechanism (capillary filling only) when its tongue was extended near its maximum during a separate feeding bout (Fig. 3). This observation sheds light on the intraoral mechanism of suction generation as we discuss further down.

### Mechanistic hypothesis testing of the tongue filling process

In addition to the fluid dynamics modeling, other lines of evidence also support the intralingual suction hypothesis. In several videos, bubbles can be seen moving inside the tongue when a bird does not make complete contact between its tongue tips and the nectar reservoir (e.g., Video S2). If capillarity was the sole mechanism used by sunbirds then no bubbles would be present inside the cylinders, as the surface tension at the distal end of the tongue would not allow bubbles to pass into the tongue. Moreover, in several instances, we observed that the nectar columns inside the tongue cylinders from the previous lick moved mouthwards before the tongue tips contacted the nectar reservoir (see trailing menisci in Videos S1, S3). This occurs whenever there is sufficient distance between the bill tip and the nectar reservoir, as the suction mechanism can begin while the tongue is being extended, regardless of its position relative to the nectar reservoir. In these instances, when the tongue is being extended, and its tips are not in contact with the nectar yet, there is a nectar-air interface at both the distal and proximal end of the tongue tubes. The distal interface creates a surface tension force pulling the nectar in the tongue towards the tip of the tongue which opposes surface tension forces applied at the unseen proximal end of the tongue cavity. The cancellation of surface tension forces means that by capillarity alone (in the absence of any external shape change) the nectar column would not rise (*40*). Inertial forces at this small scale can also be safely disregarded as the cause of the intralingual displacement of the nectar columns. Since the nectar does, in fact, move toward the mouth before the tongue contacts the reservoir, an additional force must be applied to result in the observed motion. Additionally, an amethyst sunbird *Chalcomitra amethystina* was recorded while its tongue suddenly dropped down from contact with its upper bill resulting in nectar flowing out of its oral cavity (Video S4). We hypothesize this unusual event resulted from the failure of a hermetic seal between the tongue and upper bill (see Discussion) as the bird moved its tongue too far out/down while feeding.

Lastly, in all feeding events, we observed throat motions that would correspond to displacement of the hyoid apparatus (bones supporting the tongue). This displacement would allow the controlled tongue depression inside the mouth to generate suction (as we discuss in our biomechanical hypothesis below).

## Discussion

For over two centuries, it was believed that the passive capillary force was the main mechanism for nectar feeding in birds (*1, 26, 35, 41*). Using fluid dynamic models, Kim and collaborators (2011) showed correspondence between the optimal concentration for caloric ingestion, using published data, and the predicted energy intake peak based on feeding mechanism. In their study, three bird groups – hummingbirds, honeyeaters, and sunbirds – were assumed to use capillarity, which they termed “capillary suction”, as their drinking mechanism (*35*). Since then, the capillary filling mechanism has been disproved for hummingbirds (*28, 38, 42, 43*), and for most honeyeaters—only one out of five species exhibited tongue kinematics consistent with capillarity (*39*); other avian nectarivores have not been studied until now.

Hummingbirds and most honeyeaters employ fluid trapping with their fimbriated tongue tips and an expansive filling mechanism to load up their tongue grooves with nectar; the latter mechanism is enabled by the dorso-ventral compression of their tongues upon protrusion, such that once in contact with the nectar reservoir, the elastic energy stored restores their shape drawing nectar in (*28, 38, 39*). To offload the nectar inside the bill and swallow it, they wring their tongues with their bill tips—which compresses the grooves—at the beginning of the subsequent lick (*30, 39, 44*). Our initial hypothesis was that sunbirds, with convergent traits in their feeding apparatus morphology, would also use similar techniques. However, our results were not consistent with this initial hypothesis. Unexpectedly, sunbirds do not use their bill tips to compress their tongues while feeding. In contrast, sunbirds keep their bills slightly open during the entire nectar feeding process, with the tongue reciprocating along the maxilla, keeping the mandible apparently disengaged from the drinking mechanism. Tongue kinematics, nectar flow observations, and fluid dynamics models demonstrated that a simple capillary filling mechanism cannot explain the nectar uptake patterns reported here for sunbirds. Instead, sunbirds move nectar through their tongues with shallower licks, over relatively longer immersion times, and at a slower licking rate than hummingbirds and honeyeaters (*30, 39*). The only mechanistic model that matched the collected data was the one that included a difference of pressure within the tongue cylinders.

We propose that sunbirds use active suction to generate a pressure gradient when drinking nectar. We present a biomechanical hypothesis for intralingual suction feeding, aided by cross-section diagrams of the feeding apparatus across the lick cycle, depicting the tongue motions that could generate such a pressure differential (Fig. 4). Initially, the tongue lies centered inside the slightly open bill; a ridge in the roof of the oral cavity matches the dorsal tongue shape (Fig. 4A XS 1) and would help to maintain the tongue laterally centered as it slides along the upper bill. When the tongue is extended to reach the nectar reservoir, its base is lifted, pressing against the mouth’s roof (Fig. 4B). We hypothesize that pressing the base upwards results in a partial flattening of the “U”-shaped (in cross section) proximal tongue region (Fig. 4B XS 2 and 3) that pushes out the air trapped between the tongue dorsal surface and the oral roof. The flexible tongue edges would create a hermetic seal against the maxilla producing a “suction cup” effect (Figs. 1B, 4B XS 2-3). Meanwhile, when the tongue tips make contact with the nectar reservoir, capillary forces initiate the wetting of the inner tongue spaces at the fringed tip and a rising meniscus can form via capillarity inside the tongue cylinders (which have a ventrolateral longitudinal slit). Although initially, portions of the intralingual nectar flow fit capillarity expectations, the overall meniscus advancement rate can only be explained by the addition of active suction (Fig. 3). We hypothesize that suction begins when the tongue base is depressed after being pressed against the roof of the oral cavity. Using synchronized side views, we observed a downward motion of the throat while nectar was simultaneously flowing inside the tongue cylinders, as observed in the top view (e.g., Video S1). This downward motion of the throat corresponded to the area where the hyoid bones that support the tongue base move through, thus consequently corresponding with the “U”-shaped cross-section of the tongue (Figs. 1C XS 1-2, 4B-C XS 2-3) being depressed within the oral cavity. The increasing space between the tongue and oral roof (sealed at the back by the proximal tongue edge, Fig. 1B) creates a region of lower pressure relative to the atmosphere which draws nectar through the tubular-shaped distal and middle regions of the tongue (Fig. 4C XS 3-4). It is important to note that although there are narrow gaps formed by the longitudinal slits along the tongue, the tightly curled and sometimes overlapping tongue edges could allow the tongue to work as a straw in the presence of a strong suction action. Next, the tongue is pulled proximally towards the throat and as it is retracted, it unseals from the maxilla, allowing the bird to swallow the nectar now in the proximal end of the oral cavity (Fig. 4D). The nectar column breaks near the proximal end of the semi-cylindrical section of the tongue, leaving some nectar trapped in the middle and distal portions of the tongue, where the internal lingual cylinders are thin enough for surface tension forces to prevent dislodging of the liquid inside (Fig. 4D). The tongue is now back at its approximate initial position—although it is not entirely retracted as in other nectarivores (*7, 39, 45, 46*)—but still contains some nectar within the lingual cylinders (Fig. 4E). To complete the ingestion of the entire nectar aliquot, as the lick cycle begins again, the tongue is extended and its proximal portion is pressed into the roof of the mouth before being pulled down to generate negative pressure. The nectar already in the lingual cylinders flows mouthwards because of the pressure differential at the tongue base (Fig. 4F). In some cases, this occurs even before the tongue tip makes contact with the nectar reservoir if the tongue needs to be extended far to reach it.

**Fig. 4.**
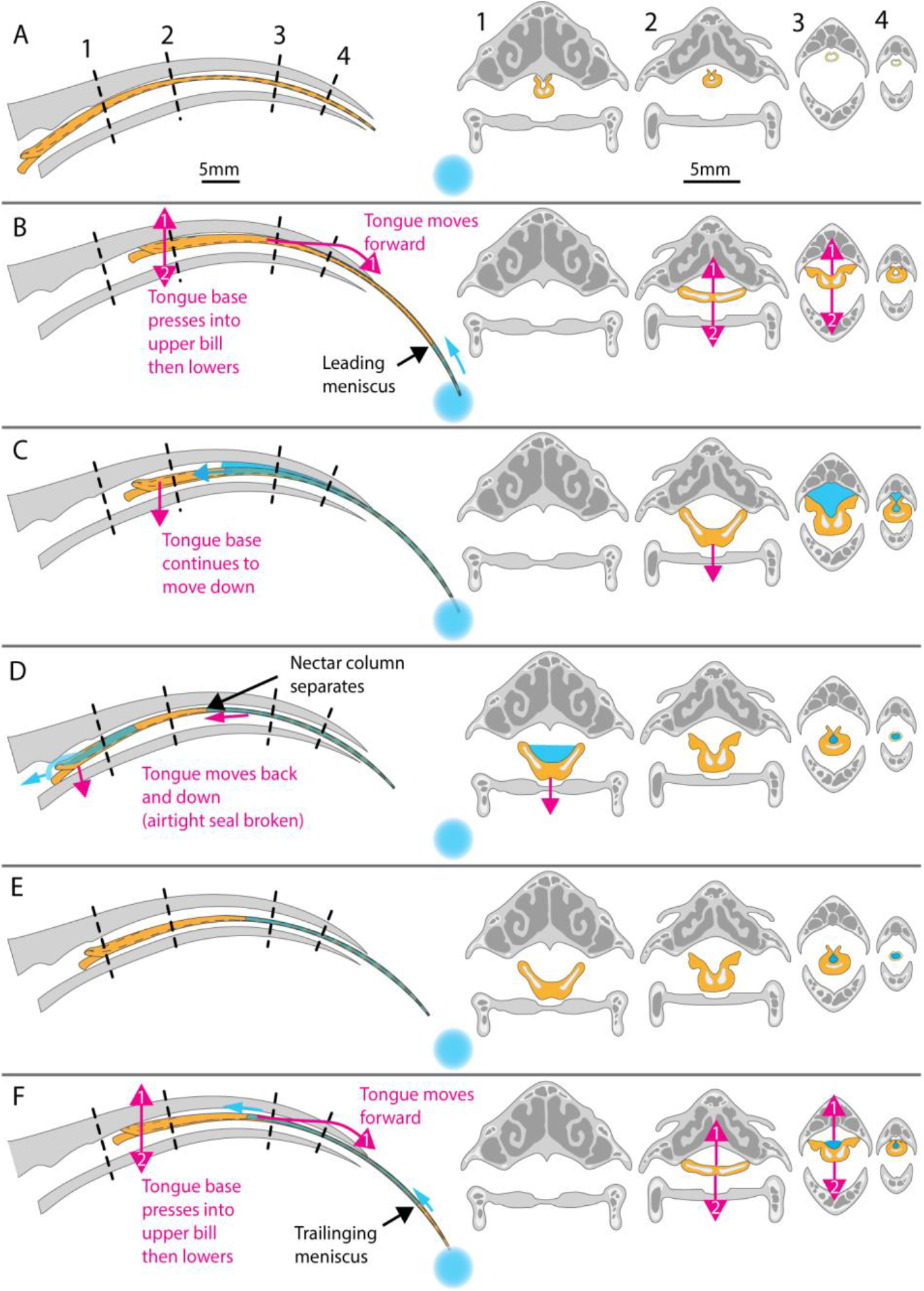
Biomechanical hypothesis of sunbird suction feeding. Sagittal cross sections of the bill and tongue of a feeding sunbird **(left)**. Coronal cross sections (XS) of the bill and tongue while feeding **(right)**. These cross sections combine anatomically accurate information from µCT scans, hypothetical motions, and deformations inferred from our external high-speed video observations. (A) Initial position of the tongue at the start of the first lick. (B) Tongue extension and base pressed into the mouth’s roof before depression. (C) The tongue continues to depress while tongue edges are sealed against the upper bill. (D) Partial retraction of the tongue, proximal portion lowered from the oral roof. The hermetic seal is released and nectar can move from the back of the tongue to the proximal oral cavity and throat. (E) The tongue has completed a lick cycle. Some nectar often remains in the tubular portion of the tongue from the previous lick cycle. (F) During re-extension for the second lick, the nectar left over from the previous cycle is drawn back into the tongue via suction as the bird begins to generate a pressure differential across the length of the tongue before the tongue tips make contact with the nectar reservoir.

In summary, several lines of evidence support the existence of a novel vertebrate feeding mechanism in sunbirds: 1) they do not immerse their tongues deep inside the nectar to use fluid trapping; 2) they do not wring their tongues with their bill tips, precluding expansive filling of the lingual cylinders; 3) the nectar flow in the tongues is closest to the fluid model for active suction, and does not decrease in flow rate over time as predicted with other models; 4) while feeding, sunbirds sometimes draw in bubbles of air into their tongues if they do not make complete contact with the nectar reservoir, meaning that capillarity alone is not responsible for the nectar flow; 5) sunbirds depress their tongues and hyoid apparatus inside their mouths simultaneously with the flow of nectar, pointing to use of tongue depression to generate a pressure differential leading to active suction; 6) as seen in an individual *C. amethystina*, when the tongue is overextended and depressed, the hermetic seal can be broken and the nectar load is lost (Video S4); and 7) as observed in an *A. malacensis* (Fig. 3), only when the tongue is extended much further than normal to make contact with the nectar reservoir the tongue base will not be in the appropriate position to generate a pressure differential and thus capillary filling is the dominant mechanism for those lick cycles. This last line of evidence shows that sunbird tongues, as shown in hummingbird tongues, have the architecture that can allow for capillary filling (*28, 42, 47*). However, this only occurs when the distance between the bill tip and nectar reservoir is farther than would be found in any flower it visits in nature. The fact that a mechanism is physically possible does not make it biologically relevant, and here we have demonstrated that suction is used in realistic conditions by sunbirds for nectar-feeding. Our findings provide significant implications for how these birds select the flowers they pollinate, as well as the coevolution with the plant rewards. This suction feeding mechanism is greatly influenced by the viscosity of the nectar offered by the plants they visit, which opens the door to predictions on the evolutionary interplay between floral nectar concentrations, bird preferences, and optimal energy extraction rates.

The mechanistically unique feeding style of sunbirds adds another interesting physical solution to the evolutionary challenge of feeding on small quantities of nectar concealed in flowers. While some insects are known to use suction through separately evolved structures (*9*– *11, 45, 46*), we revealed a previously unknown method to generate suction through a straw-like tongue in vertebrates without cheeks or lips. This discovery raises new mechanistic questions for other vertebrate and invertebrate groups that have solved the challenge of extracting small-volume and hard-to-reach liquid food with the many diverse materials, structures, and physics phenomena of which the biological world is composed.

## Supplementary Materials

### Materials and Methods

#### Experimental Design

We used high-speed cameras to capture the nectar, tongue, and bill movement of nine individuals from seven sunbird species (Table S1) while they fed on nectar from artificial flowers (described below). Sunbird species in South Africa (n = 5, Table S1) were captured in the Umgungundlovu district of KwaZulu Natal in grassland areas with aloes and foothill areas with proteas between June and August of 2022 and 2023. Experiments with these species were conducted at the University of KwaZulu-Natal Pietermaritzburg in the outdoor Animal House aviaries (Permits: DAFF Section 20, 12/11/1/5/2418(HP); EKZNW, OP 3034/2022; AREC, AREC/00004060/2022; AREC, AREC/00006760/2024). Sunbird species from Indonesia (n = 2, Table S1) were captured on Gunung Ambang and Gunung Klabat in North Sulawesi, Indonesia in July 2023. Experiments with these species were conducted in field camps established at each site (Permits: BRIN, 346/SIP/IV/FR/6/2023; IACUC, 4498-05). The artificial flower (Fig. S1) consisted of a red plastic ring attached to the end of a clear rectangular tube made of 4 glass microscope slides: one on the bottom, two glued on top of the bottom one with ∼5mm separating them, and the fourth glued on top of those to form a 5.9 mm width x 1.5 mm height rectangular tube. The apparatus was oriented horizontally and filled with artificial nectar (20% w/w sugar concentration, within range of local flower nectar). When birds fed from the artificial flower, they partially inserted their bills through the red disk into the tube and extended their tongues to feed. Two high-speed cameras (Chronos 1.4, Krontech) were used, one oriented to film the clear tube from above, and a second filming the side of the bird’s bill and head. The artificial nectar was dyed red with food coloring to help visualize the nectar moving inside of the sunbird tongues. The cameras used Nikkor 105 mm macro lenses (Nikon) and a film rate of 1000 to 4000 frames/second with 1280×1024 to 1280×240 pixels.

### Video Analysis

We analyzed the video data using dltdv8a (*48*), tracking the tongue tips, bill tips, nectar meniscus in the corolla of the artificial flower, and nectar menisci inside the tongue for multiple licks across multiple feeding bouts for each individual sunbird recorded in the experiment.

Corolla nectar meniscus is defined as the edge of the nectar in the clear prismatic chamber at roughly the midpoint of the tube’s width. The tongue nectar menisci are defined as the leading surface of nectar as it is drawn into the tongue and the trailing surface of the column of nectar as it moves through the tongue (these are found at different times in the feeding process, Video S1, S2, S3). The video data was calibrated using the known width of the nectar reservoir (5.9 mm) and the recording frame rate.

### Morphological analysis of bills and tongues

We took measurements of dissected tongues using light microscopes to input the effective internal tongue diameter into the fluid model equation. We also used micro-computed tomography (microCT) scans of an olive sunbird, *Cyanomitra olivacea*, at ∼20 micron resolution to compare morphology and measured internal tongue and oral cavity volumes to validate our light microscope measurements. The CT scanner x-ray source voltage and current was 42 kV and 560 microAmps, respectively. We analyzed the scan using the 3D Slicer software and the SlicerMorph package (*49, 50*). Transverse section images were taken along the bill and tongue at six points that showed relevant anatomical features of interest.

## Supplementary text

### Sunbird and hummingbird comparisons

Sunbird and hummingbird tongues share a similar distal morphology and notably distinct proximal morphology (Fig. S2). Both have tongues that are long, thin, keratinous structures with longitudinal grooves used to collect nectar reaching deep inside flowers. At the tip (Fig. S2, cross-section 5), both are bifurcated tubes with fringed ends. The distal section of the hummingbird tongue has two separate grooves, while that of sunbirds has bifurcated tips that fuse to one groove near the tip. In both groups, the tongues can fill with nectar in the distal to middle sections of their tongue (Fig. S2, XS4), but differ notably at the proximal portion of the tongue. Hummingbird tongues become a solid keratin structure at their longitudinal midpoint with no internal space to transport nectar within the tongue, whereas sunbird tongues unfurl laterally at the base, increasing the volume the tongue could carry (Fig. S2, XS 1 and 2). The tongue morphology of hummingbirds aids in their feeding as they need a strong, rigid structure to keep the tongue oriented properly when it is pushed between the compressed bill tips. This allows the hummingbird tongue to store elastic energy before rapidly expanding and filling its hollow portions with nectar, in a process known as elastic-filling. Sunbird tongue morphology, by contrast, consists of a continuous groove, opening the possibility of continuous flow through the entire length of the tongue.

Sunbird and hummingbird bills also show significant convergence. Both are long and narrow, often decurved, and have internal structures that may interact with the tongue and aid in nectar-feeding (Fig. S2; (*38*). It has been hypothesized for hummingbirds that the protrusions on the inner surfaces of the upper and lower bills guide the tongue while it is protruded and help compress the tongue at the bill tip to wring the tongue of the nectar it held from the previous lick and to elastically load the tongue for the next expansion (*38*). Sunbird bills also have some protrusions on the internal surfaces (Fig. S2). However, when the bill is closed the tongue is not compressed at any point along its length.

The flowers that sunbirds and hummingbirds feed from often have relatively long, narrow, and tubular corollas with nectar located at the base. However, the two bird lineages use different biophysical solutions with superficially convergent adaptations of their feeding apparatus to extract energy efficiently enough from the flowers they co-evolved with. These differences likely arose from their very different evolutionary origins: sunbirds evolved from a general passerine form, while hummingbirds evolved from a clade comprised of nocturnal birds. Comparing across taxonomic classes, we now find that both a bird clade, sunbirds, and at least one species of bat share similar nectar-feeding adaptations: both convey nectar along the length of the tongue, although by different means (*7*).

## Derivation of fluid models

The equation of motion of a liquid flowing into a vertical tube with a circular cross-section of radius R_T_ is given by the Bosanquet equation (*51, 52*):

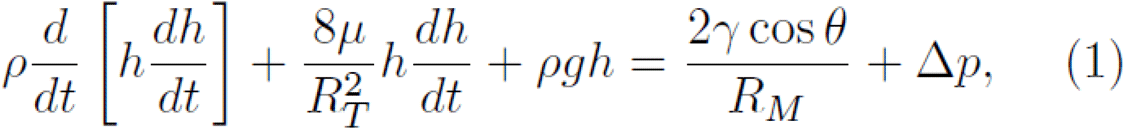

where ρ, µ and γ are the density, viscosity and surface tension of the liquid respectively. h and RM are the height and radius of curvature of the air-liquid meniscus inside the tube and θ is the contact angle between the liquid and the tube. The terms on the left-hand side of this equation represents, respectively, inertial forces, viscous forces and hydrostatic pressure and are balanced on the right-hand side by the capillary forces and an applied difference of pressure Δp. In our case, because the inner part of the tongue (tube) is wet, we assume θ = 0. In addition, because the radius RT of the tongue (≃ 0.15 mm) is much smaller than the capillary length ℓ_c_ = [γ/ρg]^1/2^ ≃ 2.7 mm, the radius of curvature of the meniscus is equal to the radius of the tube RM = R_T_.

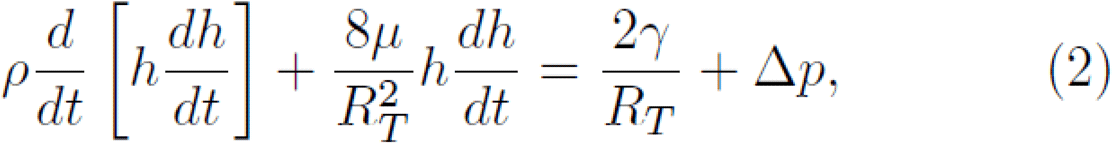

In the absence of applied pressure, this equation admits a stationary solution, i.e. dh/dt = 0, corresponding to the maximum height reachable by the meniscus, i.e. the so-called Jurin’s height: h_∞_ = 2γ/(ρgRT) ≃ 9 cm. Since, h_∞_ is significantly larger than the height reached by the meniscus in the data we analyzed, gravity can be neglected. In addition, in our case, a time-dependent applied pressure is considered which makes the role of gravity completely negligible. Therefore, Eq. (1) becomes

It is usual to neglect inertia to describe a capillary rise since its effects are noticeable only at very short time. In this case, in the absence of applied pressure, Eq. (2) becomes

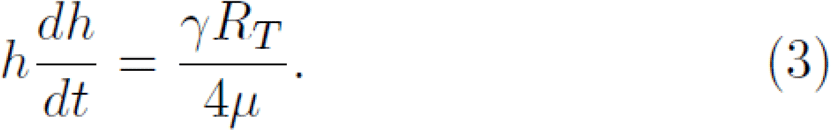

Using the initial condition h(0) = h_0_, the solution reads

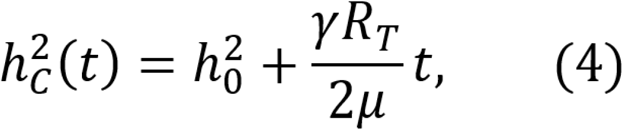

This square-root behavior does not describe well the data as shown in Fig. 3. Taking into account inertia without applied pressure, i.e. Δp = 0 in Eq. (2), together with h’(0) = 0 leads to the following solution

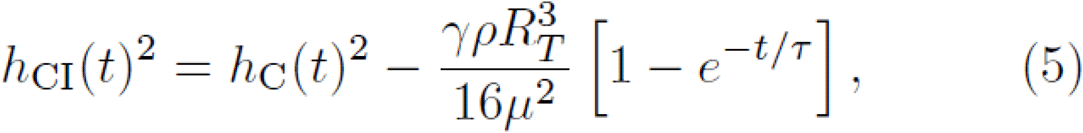

where τ = ρRT^2^ /(8*µ*) gives the typical time below which inertia dominates the viscous effects. Fig. 3 shows that adding inertia does not improve the description of the data since the curve is always below the curve without inertia as seen in Eq. (5). This indicates that the capture of nectar by sunbirds is not passive.

Finally, solving Eq. (2) with an applied pressure varying linearly in time, i.e. Δp(t) = α t, leads to

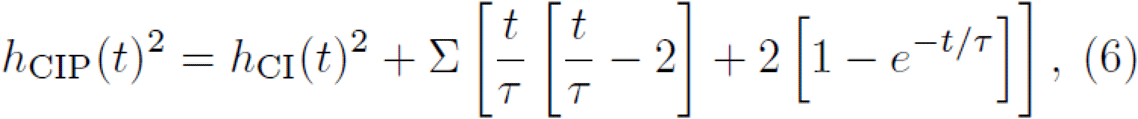

where Σ = αρ^2^RT^6^ /(8*µ*)^3^. Note that, in our case, τ is quite small (about 1.5 ms) and Eq. (6) can be simplified by expanding hCIP for t >> τ:

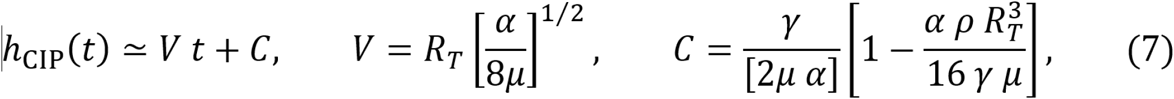

where V is the speed at which the liquid rises in the tongue and which results from the competition between the applied pressure driving the motion and the viscosity slowing down the dynamics. The constant C is a small offset resulting from the competition between all effects involved at small time t << τ (inertia, capillarity, viscosity and applied pressure) as seen by the presence of ρ, γ, µ and α in its expression.

Eqs. (4)-(6) are compared to in vivo measurements in Fig. Z with γ = 0.07 N/m, µ = 0.002 Pa s, ρ = 1085 kg/m^3^ and the values of R_T_ and α reported in Table S2 for each species. The difference of pressure applied by the birds during the time interval where the data have been measured is small compared to the atmospheric pressure.

With the values of the parameters mentioned above and reported in Table S2, V given in Eq. (7) varies between 0.4 and 0.8 m/s.

## Supplementary Figures

**Fig. S1.**
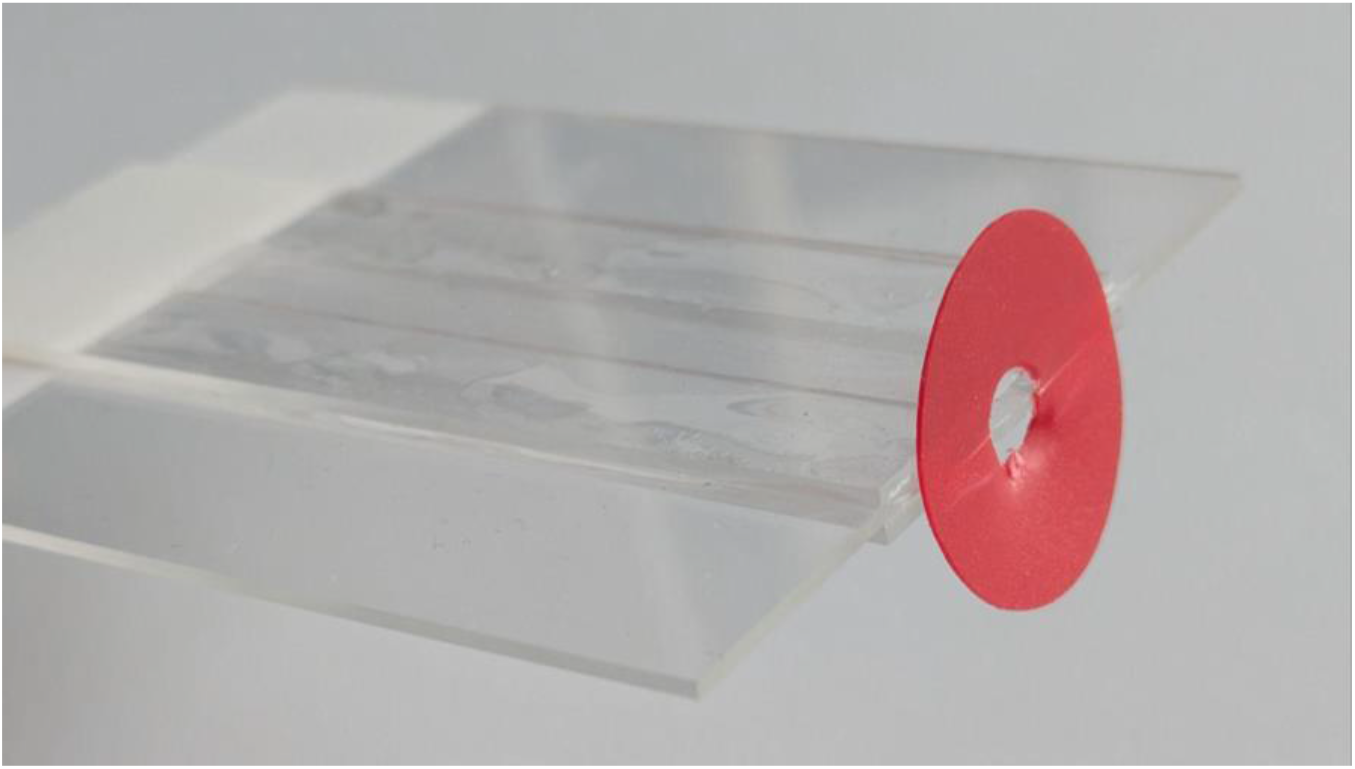
Image of the artificial flower. Made using 4 microscope slides glued together to make a 1mm x 6mm x 75mm hollow channel for nectar to be added. A red plastic ring is attached to the front to attract the bird. The apparatus is oriented horizontally for all trials to eliminate gravitational effects.

**Fig. S2.**
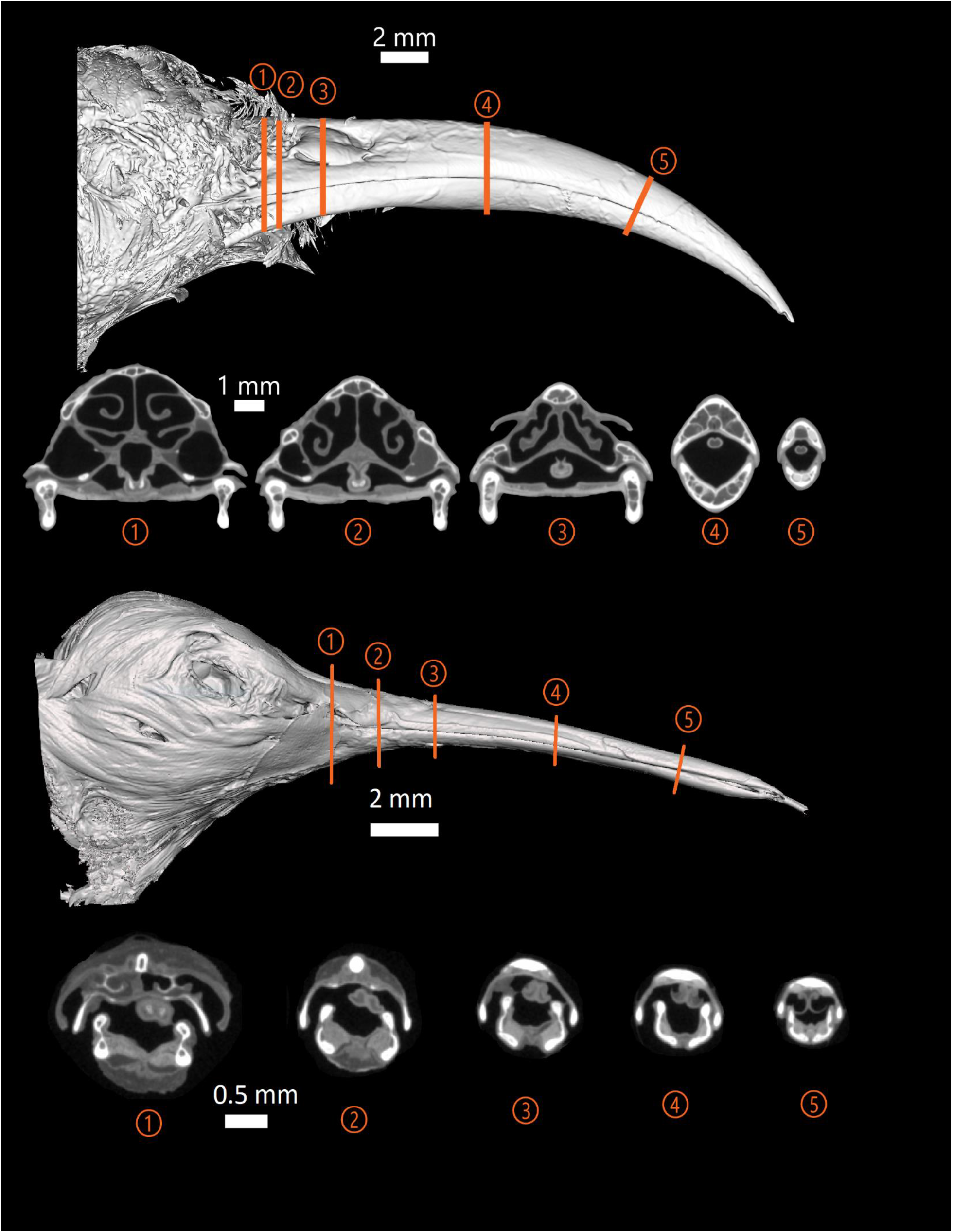
Cross-sections of sunbird (*Cyanomitra olivacea*, top) and hummingbird (*Calypte anna*, bottom) CT scans showing details of bill and tongue. Lateral views of a sunbird and hummingbird with multiple sectioning lines and corresponding CT cross-sections are shown from the proximal to distal regions of the bill and tongue. The shape of the sunbird tongue can be seen as a “U” shaped trough at the proximal end and an enclosed tube-like structure at the distal end. The hummingbird tongue is made of two solid rod-like structures at the proximal end that become two hollow tubular structures towards the distal end.

### Supplementary tables

**Table S1.**
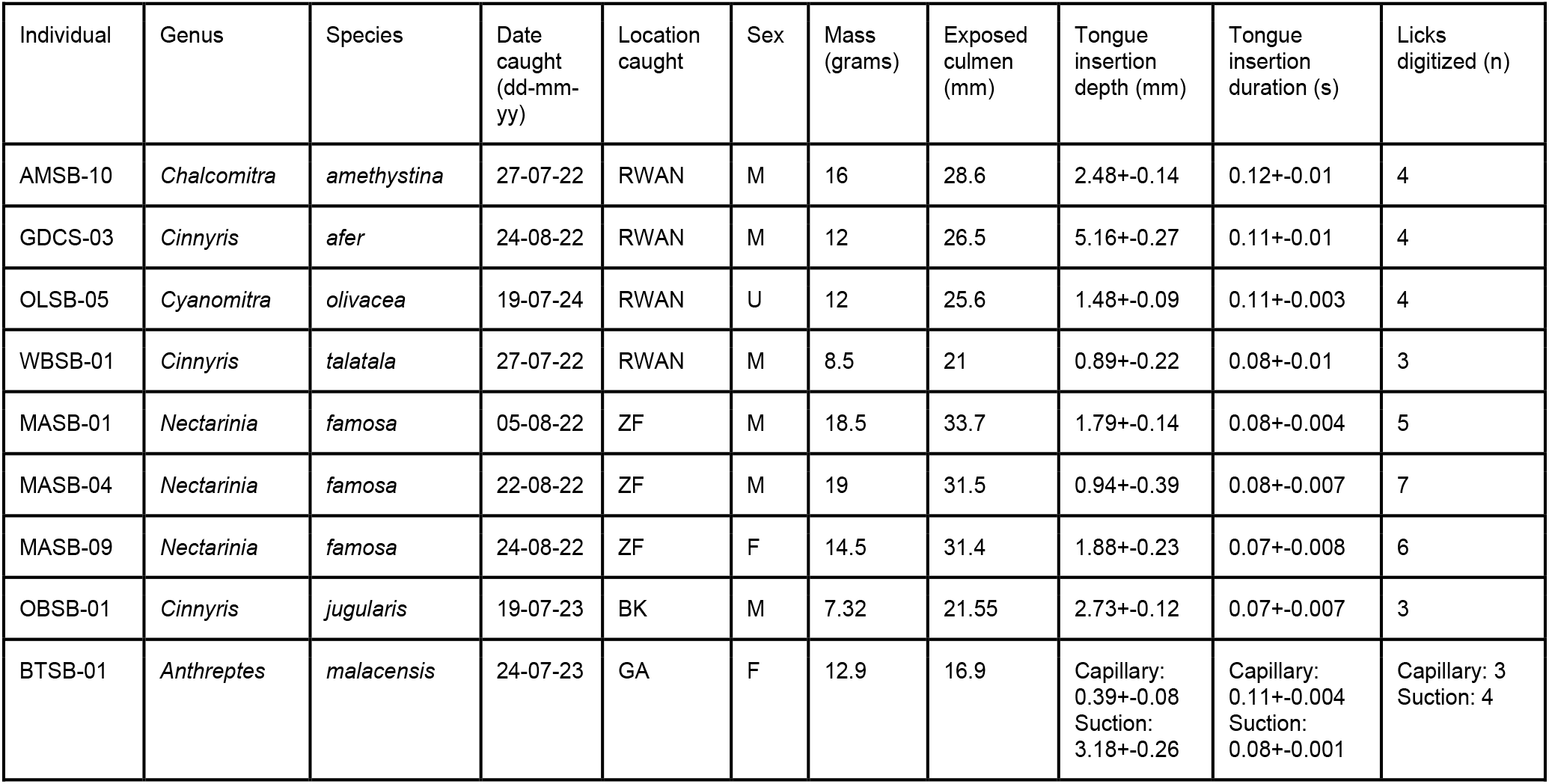
Data for individual sunbirds included in the study. Locations are RWAN: Rocky Wonder Aloe Nursery (−29.638591N, 30.491850E), ZuluFlora (−29.533766N, 30.353995E), Bongkudai (0.764167N, 124.441936E), and Gunung Klabat (1.437379N, 125.000389E). Exposed culmen is the straight line length between the bill tip and the most proximal portion of the exposed keratin on the upper bill. Tongue insertion depth is the maximum distance between the nectar reservoir meniscus and the tongue tip when submerged.

**Table S2.**
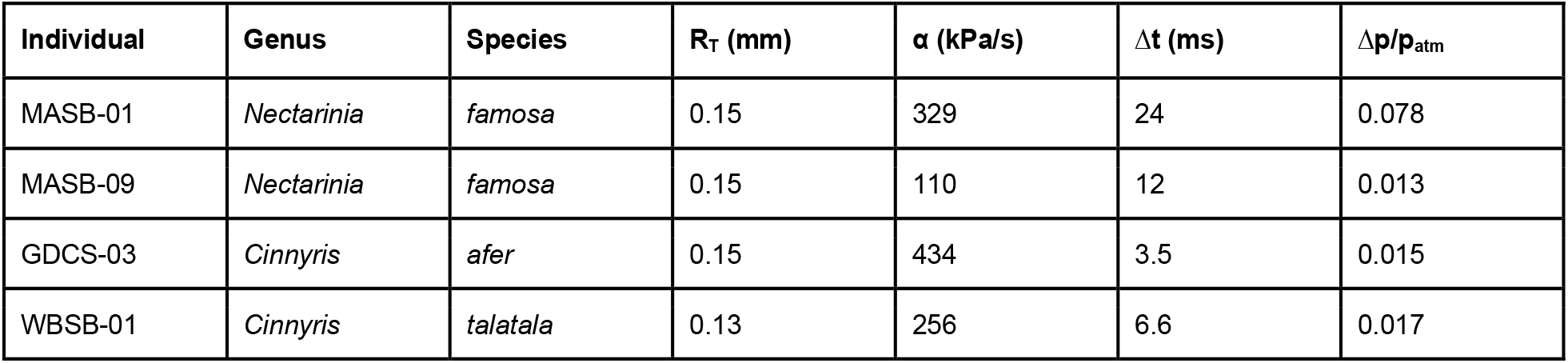
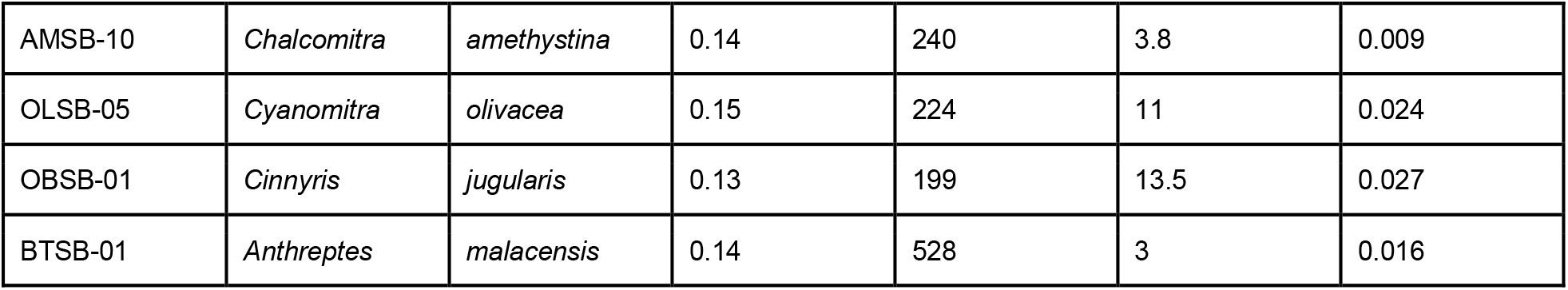
Fluid dynamics model variables. R_T_ is the effective radius of the tongue along the enclosed, tubular length. α is the pressure differential per second applied by the bird. Δt is the time period over which the bird applies the pressure differential. Δp/p_atm_ is the total applied pressure relative to the ambient atmospheric pressure.

### Videos

**Video S1.**
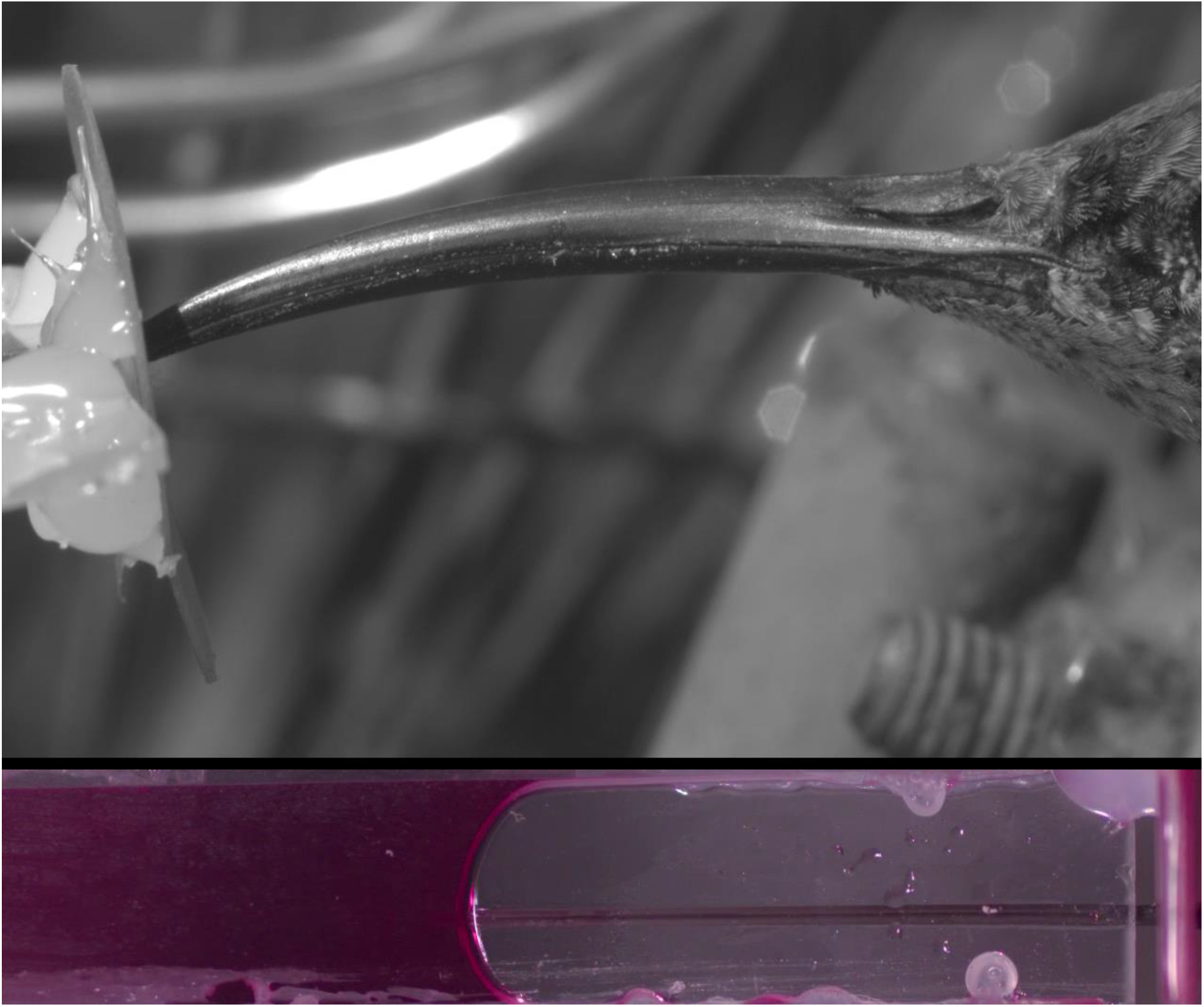
[Video_S1.mp4](https://drive.google.com/file/d/1fsDwljV-4VbnOdCiF2Cd5n6De_P0DvjR/view?usp=sharing) Top: lateral view of a malachite sunbird (*Nectarinia famosa*) feeding from the artificial flower. Bottom: dorsal view of the artificial flower (opening on the right) with the malachite sunbird’s tongue visible in the flower making contact with the red-dyed nectar.

**Video S2.**
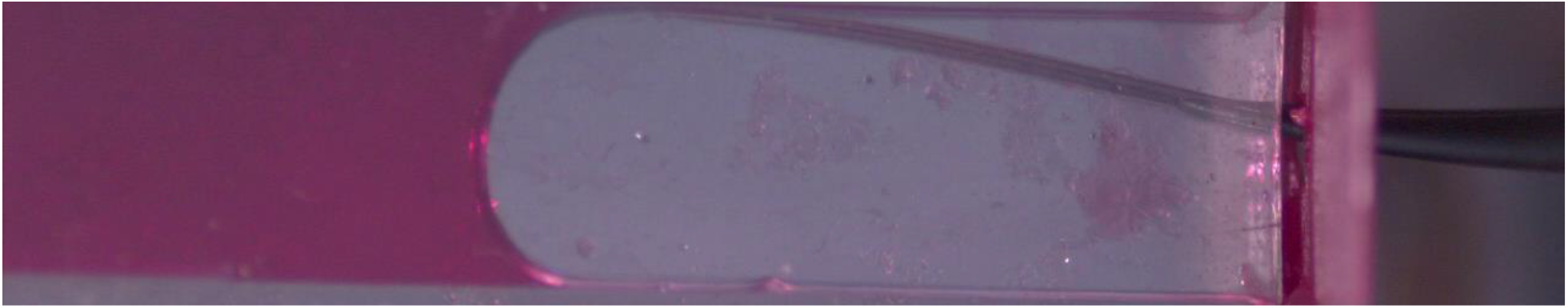
[Video_S2.mp4](https://drive.google.com/file/d/1zeHNttF8WfjlDSS0ny9PAYpaD1fpFOX0/view?usp=sharing) Dorsal view of the artificial flower with a greater double-collared sunbird *Cinnyris afer* feeding. Bubbles can be seen moving through the tongue towards the mouth (to the right) when the bird does not make complete contact between the nectar reservoir and its tongue.

**Video S3.**
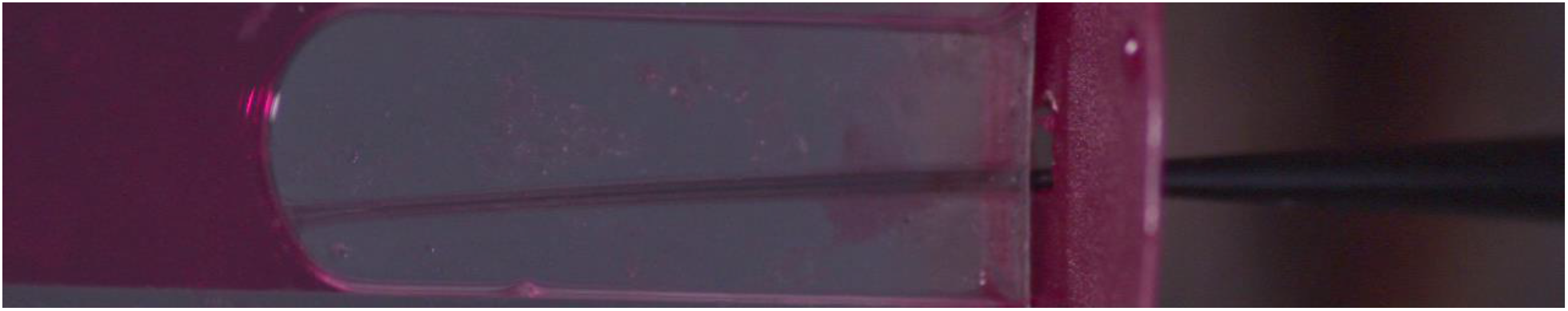
[Video_S3.mp4](https://drive.google.com/file/d/1ARrUYEjehZVeM_YcgQDKbuWpDnU94f5V/view?usp=sharing) Dorsal view of the artificial flower with a malachite sunbird *Nectarinia famosa* feeding. Nectar can be seen flowing in the tongue towards the mouth (to the right) before the tongue makes contact with the nectar reservoir.

**Video S4.**
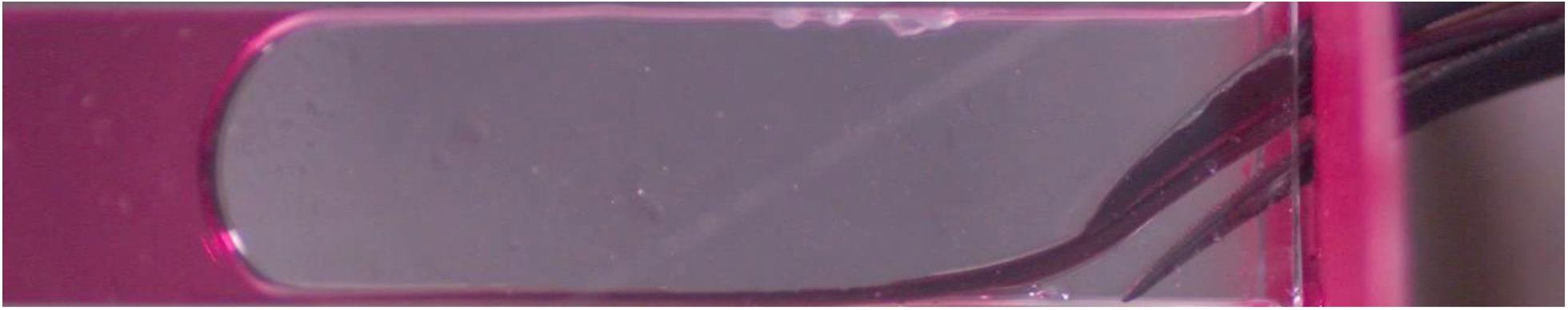
[Video_S4.mp4](https://drive.google.com/file/d/1HyXsmwFl3ZO3QB_gIe91HFGLAQ6SpbiL/view?usp=sharing) Dorsal view of the artificial flower with an amethyst sunbird *Chalcomitra amethystina* feeding. The tongue is over depressed and unseals from the upper bill and nectar can be seen escaping the mouth.

